# AN ADMIXTURE SIGNAL IN ARMENIANS AROUND THE END OF THE BRONZE AGE REVEALS WIDESPREAD POPULATION MOVEMENT ACROSS THE MIDDLE EAST

**DOI:** 10.1101/2020.06.24.168781

**Authors:** Anahit Hovhannisyan, Eppie Jones, Pierpaolo Maisano Delser, Joshua Schraiber, Anna Hakobyan, Ashot Margaryan, Peter Hrechdakian, Hovhannes Sahakyan, Lehti Saag, Zaruhi Khachatryan, Levon Yepiskoposyan, Andrea Manica

## Abstract

The Armenians, a population inhabiting the region in West Asia known as the Armenian Highland, has been argued to show a remarkable degree of population continuity since the Early Neolithic. Here we test the degree of continuity of this population as well as its plausible origin, by collating modern and ancient genomic data, and adding a number of novel contemporary genomes. We show that Armenians have indeed remained unadmixed through the Neolithic and at least until the first part of the Bronze Age, and fail to find any support for historical suggestions by Herodotus of an input from the Balkans. However, we do detect a genetic input of Sardinian-like ancestry during or just after the Middle-Late Bronze Age. A similar input at approximately the same time was detected in East Africa, suggesting large-scale movement both North and South of the Middle East. Whether such large-scale population movement was a result of climatic or cultural changes is unclear, as well as the true source of gene flow remains an open question that needs to be addressed in future ancient DNA studies.

## Introduction

The Armenian population has historically inhabited the Armenian Highland, a mountainous region stretching from the Mediterranean northeast to the South Caucasus (Fig.1, A). Geographic isolation, distinct language, a strong national and cultural identity might have been conducive to the long-term genetic isolation of Armenians from the neighboring populations [1,2,3]. Indeed, a comparison of both ancient and modern Armenian mitochondrial genomes (mtDNA), across a time span of eight thousand years, revealed a remarkably high level of matrilineal genetic continuity in the region [1]. The picture of genetic isolation is further backed up by genome-wide autosomal studies of contemporary Armenians suggesting a lack of an external genetic influx at least since the Bronze Age [2,3]. However, contemporary genomes have limited power to draw inferences on the genetic continuity through time.

**Figure 1.**
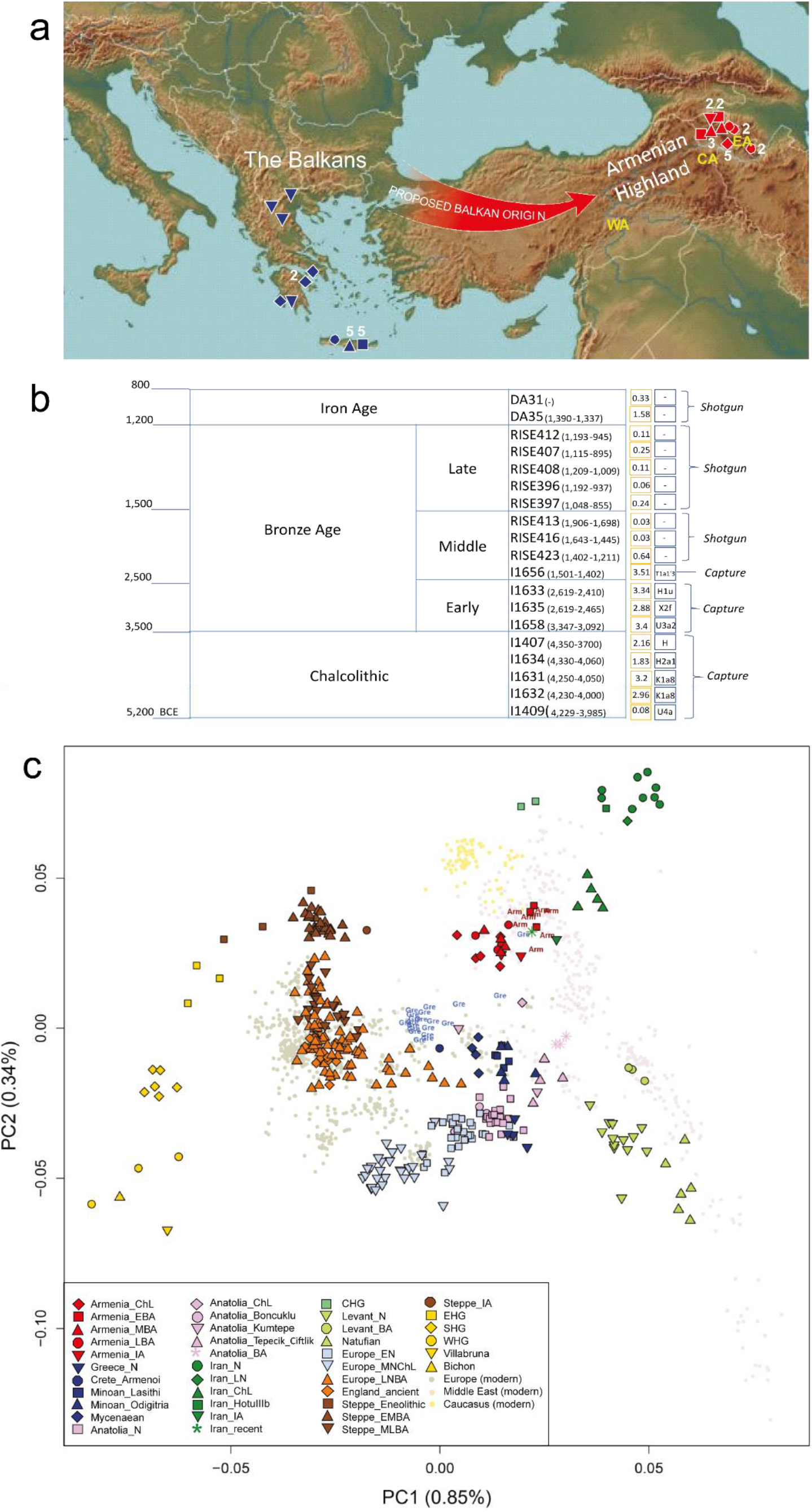
(a) Proposed route of Armenian migration according to the Balkan theory. Symbols represent populations and their shapes are described on the panel c. The numbers next to them indicate a sample size (if there is no number then the sample size is equal to one). WA, CA, EA are the abbreviations for modern Armenian samples from the western, central and eastern parts of the Armenian Highland, respectively. (b) Table with ancient Armenian samples. Radiocarbon dates (in cal BP) are shown under the sample name. Mean genome coverage is shown in yellow squares, mitochondrial haplogroups in blue squares (c) Principal component analysis (PC1 *vs* PC2). Values in parenthesis represent the percentage of variance explained by a given PC. According to the PCA, ancient samples from the Armenian Highland cluster with contemporary Armenians, while modern and ancient samples from the Balkans show distinctive clustering closer to other European populations.

The origin of Armenians is highly debated. Several theories and legends exist as to the formation of the population, though two of them prevail. According to the broadly accepted Balkan theory based on the ancient Greek historian Herodotus’ writings, the ancestors of the Armenians were Phrygian colonists who migrated to the Armenian Highland from the Balkans [4]. The conclusion was derived mainly from the fact that Armenians were armed in the Phrygian fashion when they were part of the Persian army. Common ancestry for the Armenians and the Phrygians is further suggested by some linguists, who speculate that proto-Armenian language belonged to a Thraco-Phrygian sub-group within the Indo-European language family [5]. However, an alternative view based on popular legend for the origins of Armenians suggests the local formation of the Armenian population [6]. Despite extensive excavations in the area, to date, no convincing archaeological evidence has been found to support either of these hypotheses. Furthermore, no thorough genetic study on the relationships between modern Armenians and the ancient and modern samples from the Balkans has been conducted.

The availability of ancient DNA (aDNA) from different geographical areas and time periods has brought direct evidence from the past. Through comparing ancient and modern samples, it has become possible to get detailed insights into the historical relationships between human populations and make powerful inferences on whether there is direct genetic continuity in the region or immigration by an exogenous population [7, 8]. Here, we collated all published genomic data for Armenians (both ancient and modern), as well as generated a number of whole-genome sequences of modern Armenians with all four grandparents (4GP) originating from western (n=2), central (n=2) and eastern (n=2) regions within the Armenian Highland (Fig. 1a,b). By putting together a dataset covering a broad timescale and range of ancestries, we were able to formally investigate the genetic origins of Armenians, their genetic continuity as well as check whether the Armenian population can indeed be considered as an isolate that was sheltered from the major migrations that shaped the rest of Western Eurasia.

## Results

### Relationships to modern and ancient populations: testing the Balkan theory

We first performed a principal component analysis (PCA) to assess the genetic affinities of the Armenian samples to a broad panel of ancient and modern-day populations (Fig.1c). We observed that the modern Armenians’ cluster falls in-between the genetic variation of the modern Caucasus and the Middle East, which is in accordance with their geographic location. Moreover, the cluster partly overlaps with the ancient inhabitants from the Armenian Highland, thus suggesting a degree of regional genetic continuity since the Chalcolithic (the earliest samples available for this region). In stark contrast, modern and ancient samples from the Balkans appear significantly distant from the Armenian cluster and are drawn mostly toward other European populations. To further investigate this pattern, we used a *D* statistics in the form *D*(Modern_Armenians, Ancient_Armenian_Highland; *X_Balkan*, Mbuti) to formally test whether modern Armenians received any genetic input from ancient and modern samples from the Balkans (Table 1, Supplementary Table 1). To account for attraction in ancient samples due to damage, we performed our analysis with transversions only. We did not observe any significantly positive values for the D statistic, implying that there is no Balkan-related ancestry in modern Armenians. Furthermore, we found that ancient and modern samples from the Armenian Highland form a clade (|Z| < 3) to the exclusion of ancient and modern samples from the Balkans. Thus, both PCA and *D* statistics support the distinctiveness of ancient and modern Armenian populations from any Balkan input.

**Table 1.** The *D* statistics of the form (*D*(Modern_Armenians, Ancient_Armenian_Highland; *X_Balkan*, Mbuti)) based on SNP-chip data to assess the clustering between the ancient and modern Armenian samples with respect to the ancient and modern samples from the Balkans (transversions only).

| A | B | X | Y | D | Z score | ABBA | BABA |
| --- | --- | --- | --- | --- | --- | --- | --- |
| **Modern and ancient Armenian samples form a clade ($\vert Z\vert < 3$) with respect to Balkan samples** |  |  |  |  |  |  |  |
| Armenian | Armenia_ChL | Greece_N | Mbuti | 0.0015 | 0.503 | 9615 | 9588 |
| Armenian | Armenia_ChL | Minoan_Lasithi | Mbuti | -0.0045 | -1.556 | 8839 | 8918 |
| Armenian | Armenia_ChL | Minoan_Odigitria | Mbuti | -0.0001 | -0.027 | 2377 | 2378 |
| Armenian | Armenia_ChL | Mycenaean | Mbuti | -0.0088 | -2.026 | 3535 | 3597 |
| Armenian | Armenia_ChL | Crete_Armenoi | Mbuti | -0.0079 | -0.643 | 402 | 408 |
| Armenian | Armenia_EBA | Greece_N | Mbuti | -0.0003 | -0.092 | 9333 | 9338 |
| Armenian | Armenia_EBA | Minoan_Lasithi | Mbuti | -0.01 | -3.106 | 8581 | 8754 |
| Armenian | Armenia_EBA | Minoan_Odigitria | Mbuti | 0.003 | 0.503 | 2341 | 2327 |
| Armenian | Armenia_EBA | Mycenaean | Mbuti | -0.0077 | -1.667 | 3433 | 3486 |
| Armenian | Armenia_EBA | Crete_Armenoi | Mbuti | -0.0102 | -0.703 | 400 | 408 |
| Armenian | Armenia_MBA | Greece_N | Mbuti | 0.0056 | 1.357 | 8513 | 8419 |
| Armenian | Armenia_MBA | Minoan_Lasithi | Mbuti | -0.0023 | -0.538 | 7908 | 7945 |
| Armenian | Armenia_MBA | Minoan_Odigitria | Mbuti | 0.0051 | 0.602 | 2208 | 2186 |
| Armenian | Armenia_MBA | Mycenaean | Mbuti | 0.0076 | 1.28 | 3169 | 3122 |
| Armenian | Armenia_MBA | Crete_Armenoi | Mbuti | -0.0203 | -1.02 | 380 | 396 |
| Armenian | Armenia_LBA | Greece_N | Mbuti | -0.0012 | -0.254 | 4848 | 4860 |
| Armenian | Armenia_LBA | Minoan_Lasithi | Mbuti | 0.004 | 0.731 | 4503 | 4467 |
| Armenian | Armenia_LBA | Minoan_Odigitria | Mbuti | 0.0061 | 0.545 | 1243 | 1228 |
| Armenian | Armenia_LBA | Mycenaean | Mbuti | -0.0035 | -0.433 | 1826 | 1838 |
| Armenian | Armenia_LBA | Crete_Armenoi | Mbuti | -0.0004 | -0.014 | 233 | 234 |
| Armenian | Armenia_IA | Greece_N | Mbuti | -0.0015 | -0.374 | 9433 | 9463 |
| Armenian | Armenia_IA | Minoan_Lasithi | Mbuti | 0.0035 | 0.801 | 8368 | 8310 |
| Armenian | Armenia_IA | Minoan_Odigitria | Mbuti | -0.0079 | -0.9 | 2154 | 2189 |
| Armenian | Armenia_IA | Mycenaean | Mbuti | -0.0074 | -1.183 | 3355 | 3405 |
| Armenian | Armenia_IA | Crete_Armenoi | Mbuti | 0.0262 | 1.244 | 381 | 362 |
| Armenian | Armenia_ChL | Greek | Mbuti | 0.0001 | 0.044 | 5052 | 5051 |
| Armenian | Armenia_EBA | Greek | Mbuti | 0.0043 | 1.542 | 4936 | 4893 |
| Armenian | Armenia_MBA | Greek | Mbuti | 0.0002 | 0.068 | 4494 | 4492 |
| Armenian | Armenia_LBA | Greek | Mbuti | 0.0014 | 0.311 | 2629 | 2622 |
| Armenian | Armenia_IA | Greek | Mbuti | 0.0006 | 0.153 | 4828 | 4822 |

### Insights into the Armenian regional continuity

We next used a maximum-likelihood approach [9] to test for continuity of ancient and modern populations of the Armenian Highland. This model assumes a scenario where changes in allele frequency from the common ancestor forward in time are explained through drift alone. We performed our analysis on capture and shotgun ancient data, separately. The test suggests a very recent drift time in modern Armenians since the common ancestor with the ancient samples, thus supporting the close relationship between ancient and modern inhabitants of this region. However, the model rejects the regional continuity hypothesis, since the ancient populations appear to have significantly larger drift times (Supplementary Table 2). This result is compatible with a scenario of admixture into modern Armenians from an outside population, which diverged earlier than the split time between the modern and ancient inhabitants of the region. As a consequence, this admixture would have increased heterozygosity in modern Armenians and thus it would exaggerate the drift time in ancient Armenians (since no genetic influx is allowed according to the model), eventually resulting in the rejection of the regional continuity. We then investigated corresponding signatures of gene flow into the Armenian Highland by using *D* statistics in the form *D*(Modern_Armenians, Ancient_Armenian_Highland; *X_ancient_population*, Mbuti). We find that a number of Bronze Age samples from the region break the clade of ancient and modern Armenian (Table 2, Supplementary Table 3); in all cases, the Bronze Age samples from the region are more closely associated with the ancient Armenian (Z<−3), suggesting an influx into modern Armenians after the end of the Bronze Age from an unknown source which is not well represented by any of the available ancient samples.

**Table 2.** The *D* statistics of the form (*D*(Modern_Armenians, Ancient_Armenian_Highland; *X_ancient_population*, Mbuti)) based on SNP-chip data to assess the clustering between the ancient and modern Armenian samples with respect to the ancient Eurasian populations (transversions only). Values with |Z| score > 3 are in bold.

| A | B | X | Y | D | Z score | ABBA | BABA |
| --- | --- | --- | --- | --- | --- | --- | --- |
| **There was a genetic influx to the Armenian Highland region after the Late Bronze/ Iron Age** |  |  |  |  |  |  |  |
| Armenian | Armenia_ChL | Steppe_Eneolithic | Mbuti | -0.0234 | **-5.715** | 6605 | 6922 |
| Armenian | Armenia_EBA | Steppe_Eneolithic | Mbuti | -0.0104 | -2.518 | 6552 | 6689 |
| Armenian | Armenia_MBA | Steppe_Eneolithic | Mbuti | -0.0121 | -2.237 | 6044 | 6192 |
| Armenian | Armenia_LBA | Steppe_Eneolithic | Mbuti | -0.0182 | -2.633 | 3350 | 3475 |
| Armenian | Armenia_IA | Steppe_Eneolithic | Mbuti | -0.0082 | -1.468 | 6196 | 6298 |
| Armenian | Armenia_ChL | Steppe_EMBA | Mbuti | -0.0111 | **-4.567** | 9578 | 9794 |
| Armenian | Armenia_EBA | Steppe_EMBA | Mbuti | -0.0115 | **-4.398** | 9308 | 9524 |
| Armenian | Armenia_MBA | Steppe_EMBA | Mbuti | -0.0137 | **-4.055** | 8442 | 8677 |
| Armenian | Armenia_LBA | Steppe_EMBA | Mbuti | -0.0191 | **-4.73** | 4811 | 4998 |
| Armenian | Armenia_IA | Steppe_EMBA | Mbuti | -0.0169 | **-4.808** | 9357 | 9678 |
| Armenian | Armenia_ChL | Steppe_MLBA | Mbuti | -0.0095 | **-4.117** | 9595 | 9778 |
| Armenian | Armenia_EBA | Steppe_MLBA | Mbuti | -0.0086 | **-3.299** | 9327 | 9489 |
| Armenian | Armenia_MBA | Steppe_MLBA | Mbuti | -0.0105 | **-3.121** | 8467 | 8645 |
| Armenian | Armenia_LBA | Steppe_MLBA | Mbuti | -0.0123 | -2.884 | 4845 | 4966 |
| Armenian | Armenia_IA | Steppe_MLBA | Mbuti | -0.0128 | **-3.676** | 9401 | 9644 |
| Armenian | Armenia_ChL | Steppe_IA | Mbuti | -0.0087 | -2.329 | 7940 | 8080 |
| Armenian | Armenia_EBA | Steppe_IA | Mbuti | -0.0086 | -2.105 | 7762 | 7897 |
| Armenian | Armenia_MBA | Steppe_IA | Mbuti | -0.0122 | -2.251 | 7112 | 7288 |
| Armenian | Armenia_LBA | Steppe_IA | Mbuti | 0.0041 | 0.598 | 4032 | 3999 |
| Armenian | Armenia_IA | Steppe_IA | Mbuti | -0.0157 | -2.798 | 7387 | 7623 |
| Armenian | Armenia_ChL | Anatolia_N | Mbuti | -0.0082 | **-3.585** | 9648 | 9808 |
| Armenian | Armenia_EBA | Anatolia_N | Mbuti | -0.0052 | -2.049 | 9399 | 9498 |
| Armenian | Armenia_MBA | Anatolia_N | Mbuti | 0.0031 | 0.916 | 8611 | 8558 |
| Armenian | Armenia_LBA | Anatolia_N | Mbuti | -0.0002 | -0.044 | 4905 | 4907 |
| Armenian | Armenia_IA | Anatolia_N | Mbuti | 0.0009 | 0.28 | 9566 | 9548 |
| Armenian | Armenia_ChL | Anatolia_ChL | Mbuti | -0.0013 | -0.318 | 5840 | 5855 |
| Armenian | Armenia_EBA | Anatolia_ChL | Mbuti | -0.0123 | -2.731 | 5701 | 5843 |
| Armenian | Armenia_MBA | Anatolia_ChL | Mbuti | 0.0008 | 0.125 | 5448 | 5440 |
| Armenian | Armenia_LBA | Anatolia_ChL | Mbuti | -0.0048 | -0.603 | 3113 | 3143 |
| Armenian | Armenia_IA | Anatolia_ChL | Mbuti | -0.0037 | -0.602 | 5301 | 5340 |
| Armenian | Armenia_ChL | Anatolia_BA | Mbuti | -0.0079 | -2.361 | 7381 | 7499 |
| Armenian | Armenia_EBA | Anatolia_BA | Mbuti | -0.0119 | **-3.142** | 7239 | 7413 |
| Armenian | Armenia_MBA | Anatolia_BA | Mbuti | -0.0034 | -0.693 | 6800 | 6847 |
| Armenian | Armenia_LBA | Anatolia_BA | Mbuti | 0.0041 | 0.67 | 3908 | 3876 |
| Armenian | Armenia_IA | Anatolia_BA | Mbuti | -0.0001 | -0.018 | 6775 | 6776 |
| Armenian | Armenia_ChL | Levant_N | Mbuti | -0.0105 | **-3.242** | 7414 | 7571 |
| Armenian | Armenia_EBA | Levant_N | Mbuti | -0.0095 | -2.622 | 7278 | 7418 |
| Armenian | Armenia_MBA | Levant_N | Mbuti | -0.0054 | -1.098 | 6761 | 6834 |
| Armenian | Armenia_LBA | Levant_N | Mbuti | -0.0016 | -0.271 | 3915 | 3928 |
| Armenian | Armenia_IA | Levant_N | Mbuti | -0.0009 | -0.188 | 6895 | 6908 |
| Armenian | Armenia_ChL | Levant_BA | Mbuti | -0.0084 | -2.742 | 7783 | 7915 |
| Armenian | Armenia_EBA | Levant_BA | Mbuti | -0.013 | **-3.894** | 7601 | 7802 |
| Armenian | Armenia_MBA | Levant_BA | Mbuti | 0.0013 | 0.276 | 7144 | 7126 |
| Armenian | Armenia_LBA | Levant_BA | Mbuti | -0.0008 | -0.149 | 4103 | 4110 |
| Armenian | Armenia_IA | Levant_BA | Mbuti | -0.0025 | -0.552 | 7199 | 7235 |
| Armenian | Armenia_ChL | Iran_N | Mbuti | -0.0021 | -0.776 | 9487 | 9526 |
| Armenian | Armenia_EBA | Iran_N | Mbuti | -0.0097 | **-3.308** | 9176 | 9356 |
| Armenian | Armenia_MBA | Iran_N | Mbuti | -0.0074 | -1.904 | 8354 | 8478 |
| Armenian | Armenia_LBA | Iran_N | Mbuti | -0.0075 | -1.586 | 4784 | 4856 |
| Armenian | Armenia_IA | Iran_N | Mbuti | -0.0104 | -2.835 | 9278 | 9473 |

### Identifying the source and time of admixture

In order to determine the nature of the genetic influx, we calculated *D* statistics in the form *D*(Modern_Armenians, Ancient_Armenian_Highland; *X_modern_population*, Mbuti) to check whether some modern groups break the clade between modern Armenians and ancient samples from the region. We did not find any significant *D* values when using transversions only, though we were able to detect a suggestive signal (Z>2) for introgression from a Sardinian-like source after Early Bronze Age (Table 3). We then boosted our power by increasing the number of SNPs by including also transitions in our analysis. Using all SNPs, we found significant *D* values for a genetic influx into modern Armenians from a Sardinian-like source when using Early and Late Bronze Age Armenian samples as our ancient reference (Supplementary Table 4). However, the result for the Late Bronze Age samples should be treated with caution, as these samples were not UDG treated and thus using transitions might lead to artifacts arising from ancient DNA damage. The result for the Early Bronze Age, on the other hand, is much more reliable as these samples were UDG treated.

**Table 3.** The *D* statistics of the form (*D*(Modern_Armenians, Ancient_Armenian_Highland; *X_modern_population*, Mbuti)) based on SNP-chip data. (transversions only).

| A | B | X | Y | D | Z score | ABBA | BABA |
| --- | --- | --- | --- | --- | --- | --- | --- |
| Armenian | Armenia_ChL | Sardinian | Mbuti | 0.0015 | 0.586 | 5066 | 5051 |
| Armenian | Armenia_ChL | Sicilian | Mbuti | 0.0021 | 0.84 | 5052 | 5031 |
| Armenian | Armenia_ChL | Italian_North | Mbuti | 0.0000 | 0.007 | 5060 | 5060 |
| Armenian | Armenia_EBA | Sardinian | Mbuti | 0.0066 | 2.286 | 4954 | 4888 |
| Armenian | Armenia_EBA | Sicilian | Mbuti | 0.0062 | 2.178 | 4933 | 4872 |
| Armenian | Armenia_EBA | Italian_North | Mbuti | 0.0043 | 1.5 | 4943 | 4901 |
| Armenian | Armenia_MBA | Sardinian | Mbuti | 0.0021 | 0.565 | 4505 | 4486 |
| Armenian | Armenia_MBA | Sicilian | Mbuti | -0.0001 | -0.018 | 4482 | 4483 |
| Armenian | Armenia_MBA | Italian_North | Mbuti | 0.0003 | 0.071 | 4499 | 4497 |
| Armenian | Armenia_LBA | Sardinian | Mbuti | 0.005 | 1.095 | 2639 | 2612 |
| Armenian | Armenia_LBA | Sicilian | Mbuti | 0.0036 | 0.812 | 2628 | 2610 |
| Armenian | Armenia_LBA | Italian_North | Mbuti | 0.0022 | 0.502 | 2634 | 2623 |
| Armenian | Armenia_IA | Sardinian | Mbuti | 0.0033 | 0.856 | 4845 | 4813 |
| Armenian | Armenia_IA | Sicilian | Mbuti | 0.0028 | 0.758 | 4825 | 4798 |
| Armenian | Armenia_IA | Italian_North | Mbuti | 0 | -0.002 | 4834 | 4834 |

Among modern-day populations, Sardinians have the highest affinity to early European farmers, and often act as a proxy for that ancestry in population genetic analyses [10, 11]. We assume that the migration more likely came from the Middle East, rather than a relatively isolated Mediterranean island. However, we were not able to find any detectable signal of gene flow from an ancient Anatolian farmer-like source (Table 2, Supplementary Table 3). According to recent studies, 38-44% of the ancestry of modern Sardinians is derived from an Iranian, Steppe and North-African-related source [12]. Likewise, we did not reveal any Iranian-related ancestry in modern Armenians that may have altered them from their regional ancestors (Table 2, Supplementary Table 3). Thus, it is more likely that the true source of gene flow to the Armenian Highland is yet unsampled, and that the time span between the date of mixture (i.e. at some point during/after the Bronze age) and all available ancient individuals of the Middle Eastern region (which are much older) is too large for the latter to act as representative sources.

Finally, we attempted to date the genetic influx based on the patterns of linkage disequilibrium with ALDER. We had to run a one population test (modern Armenians as the target with an input from Sardinians), which has limited power, and so we augmented our sample sizes to 99 Armenians and 23 Sardinians with 709,810 SNPs. We were able to detect a significant signal of admixture with a Sardinian-like source in Armenians, with an estimated contribution of 28.0 +/− 3.8%. The timing of the admixture obtained from ALDER suggests a relatively old event (the best signal corresponds to 172.56 +/− 17.23 generations ago), which is closer to the end of the Early Bronze Age in Armenia. However, from our the aDNA data, we know that the admixture might have occurred even later, potentially after the end of the Bronze Age (approximately 3000 ya). Such a scenario is at the limit of the dating estimated by ALDER – the end of the Bronze Age is between 3 to 4 standard deviations from the best estimate.

## Discussion

Our study, based on a combined dataset of modern and ancient genomes from the Armenian Highland, revealed a strikingly high level of regional genetic continuity for well over six thousand years, with only one detectable input from a Sardinian-like source during the Late Bronze Age or after. A recent study also suggest a similar level of stability up to the Bronze Age in the South Caucasus (there was no test for later inputs as this study used only aDNA) [13]. This pattern stands in contrast to most other Western Eurasian populations, which have undergone multiple large influxes [10,14]. A high level of continuity sets the Armenian population apart even in comparison to the Sardinians, who have long been considered as a genetic isolate in the region since the Neolithic, but recent studies have shown that the island received numerous genetic inputs after the Bronze Age [12].

We focused on solving a long-standing puzzle regarding Armenians’ genetic roots. Although the Balkan hypothesis has long been considered the most plausible narrative on the origin of Armenians, our results strongly reject it, showing that modern Armenians are genetically distinct from both the ancient and present-day populations from the Balkans. On the contrary, we confirmed the pattern of genetic affinity between the modern and ancient inhabitants of the Armenian Highland since the Chalcolithic, which was initially identified in previous studies [15–17].

However, we found that the genetic continuity in the Armenian Highland was disrupted during the second half/after the end of the Bronze Age by an input from a Sardinian-like source. The time scale for this input, as well as its source, coincide with a similar event detected in East Africa [18,19], suggesting large scale population movements from the Middle East going North and South. Our analyses failed to find a good source for this expansion among the available aDNA samples.

Our results on the source population and date for mixture are mostly in line with the studies on Armenians conducted so far. An elevated Sardinian-like ancestry in Armenians was also detected in the previous study [2] based on modern genomes, and the gene flow rate was estimated as 29%, which is quite comparable with our findings. Our study including aDNA samples enhances the power of admixture tests by providing proper sources for population ancestry. As such, we could not find any evidence for mixtures of multiple populations during the time period of 3,000–2,000 BCE. In contrast, we found a signal for a single admixture event from a Sardinian-like source which happened after the Early Bronze Age, and possibly after the end of the Bronze Age (but the lack of UDG treated samples for the latter part of this period in Armenia warrants some caution).

## Conclusions

Thus, we conclude that there was large-scale movement across the Middle East towards the end or after the Bronze Age. From the genetic point of view, this movement has led to an essential input of Neolithic ancestry in the populations inhabiting the region. In particular, as a result of its movement northward, the population of the Armenian Highland has become genetically altered from its regional ancestors. While Sardinians are the best proxy for this gene flow, questions like where it comes from and what was a trigger for such a widespread migration wave are still unanswered. To investigate this further, further studies on the complex demographic processes of the region should be conducted with the incorporation of more ancient and modern data.

## Materials and Methods

### Sample collection and sequencing

Blood samples were collected from unrelated individuals with all four grandparents (4GP) originating from the western (n=2), central (n=2) and eastern (n=2) parts of the Armenian Highland. This study was approved by the Ethics Committee of the Institute of Molecular Biology NAS RA (IRB #00004079). Armenian subjects were informed about the aim of this study and gave their consent to participate.

DNA samples were extracted at the Institute of Molecular Biology NAS RA using a modification of the salting out procedure [20]. The DNA was normalized (~50 ng/μl) and sent for sequencing (Macrogen Company, South Korea) at high coverage (~30x) using paired-end whole genome sequencing on the Illumina HiSeq X Ten.

### Data processing and alignment

Adapter sequences were trimmed from the ends of reads using *trimadap* [21]. Sequences were aligned to the hs37d5 build of the human reference genome using Burrows-Wheeler Aligner (BWA) version 0.7.16 [22] and clonal reads were removed with *samblaster* [23]. Variant calling was performed using Genome Analysis Toolkit (GATK) version 2.5.2 [24].

### Merging with ancient and modern reference dataset

We merged our six modern Armenian genomes with the previously published two Armenian (4GP) whole genomes from the Simons project [25]. Fastq data were aligned and genotyped with the same pipeline described above. We used PLINK 1.09 [26] to combine our modern Armenian samples with calls from modern populations in the Human Origins (HO) dataset [10] and ancient samples from Lazaridis et al., 2017 [14] which also includes genotype calls for previously published ancient individuals [15, 16, 27–29]. We downloaded FASTQ files of Iron Age samples from the Armenian Highland [30], aligned them to the hs37d5 build of the human reference genome using BWA and duplicate reads were removed using SAMtools version 0.1.19 [31]. Indels were realigned using GATK RealignerTargetCreator and IndelRealigner [24]. Sequences with a mapping quality of ≥ 30 were retained with SAMtools [31]. Pileupcaller was used to sample alleles from low-coverage sequenced Iron Age Armenian genomes [32]. The merging with other dataset led to 1,054,527 SNPs when modern Armenians and ancient samples were analyzed, and to 591,558 when modern samples from the HO dataset were also included.

### Population genomic analyses

Principal component analysis (PCA) was performed with a broad panel of modern and ancient samples using the *smartpca* software from Eigensoft 7.2.0 [33] with the outlier removal option off. All ancient samples were projected onto the principal components by using the options lsqproject:YES.

*D*-statistics on SNP array data were computed using the qpDstat program in the ADMIXTOOLS package [34]. Statistics were considered significant if Z-score was greater than 3 corresponding to a P-value <0.001. To account for the potential effect of ancient DNA damage, we repeated our analyses both with and without transition substitution sites [35].

A formal continuity test was performed using the method described in Schraiber 2018 [9]. Prior to the analysis, we fitted alpha and beta priors to the discrete reference allele frequencies. Files of ancestral alleles were downloaded from the following source: ftp://ftp.1000genomes.ebi.ac.uk/vol1/ftp/phase1/analysis_results/supporting/ancestral_alignments/. We restricted the analyses to sites with high-confidence calls where the ancestral state is supported by all sequence comparisons. We also did not include alleles with the frequency 0 or 1 in the modern population. Thus, for tests of population continuity with capture data, we were able to use 685,797 SNPs. For shotgun data, prior to running the analyses, we performed LD pruning with the option–indep-pairwise 200 25 0.4 and further random subsampling of half of the remaining SNPs. This resulted in 1,826,787 SNPs available in the test of population continuity.

To date the time of admixture, we used ALDER [36] (version 1.03) with mindis 0.005. We processed FASTQ data on 23 Sardinian samples from Human Genome Diversity Project (HGDP)-CEPH panel [37] with the similar pipeline as for modern Armenian genomes. We combined those with data on 99 Armenians from Haber et al [2] genotyped for 710,870 snps.

## Data accessibility

Raw reads and BAM files are available for download through the accession number (will be available upon the publication).

## Authors’ contributions

A.M., A.H. and L.Y. conceived and designed the study. L.Y. and Z.K. collected the data. A.H. performed the bulk of the bioinformatics and statistical analyses under the supervision of A.M. as well as prepared all the figures. E.J., P.M.D., J.S., A.Hak., As.M., H.S., and L.S. contributed to the analyses. A.H. and A.M. wrote the manuscript; all authors contributed to its editing and approved the final version.

## Acknowledgements

We thank the volunteers for their support for this research.

## Footnotes

Electronic supplementary material is available online here ()

