## supplementary material for "AN ADMIXTURE SIGNAL IN ARMENIANS AROUND THE END OF THE BRONZE AGE REVEALS WIDESPREAD POPULATION MOVEMENT ACROSS THE MIDDLE EAST"

### Supplementary Materials

| A | B | X | Y | D | Z score | ABBA | BABA |
|---|---|---|---|---|---|---|---|
| Armenian | Armenia_ChL | Greece_N | Mbuti | -0.001 | -0.539 | 51766 | 51871 |
| Armenian | Armenia_ChL | Minoan_Lasithi | Mbuti | -0.0062 | **-3.275** | 47591 | 48186 |
| Armenian | Armenia_ChL | Minoan_Odigitria | Mbuti | -0.0086 | **-3.064** | 12924 | 13149 |
| Armenian | Armenia_ChL | Mycenaean | Mbuti | -0.0091 | **-3.865** | 19093 | 19442 |
| Armenian | Armenia_ChL | Crete_Armenoi | Mbuti | -0.0001 | -0.009 | 2276 | 2277 |
| Armenian | Armenia_EBA | Greece_N | Mbuti | 0.0018 | 0.837 | 50657 | 50478 |
| Armenian | Armenia_EBA | Minoan_Lasithi | Mbuti | -0.0049 | -2.334 | 46744 | 47200 |
| Armenian | Armenia_EBA | Minoan_Odigitria | Mbuti | -0.002 | -0.642 | 12835 | 12886 |
| Armenian | Armenia_EBA | Mycenaean | Mbuti | -0.0087 | **-3.347** | 18661 | 18991 |
| Armenian | Armenia_EBA | Crete_Armenoi | Mbuti | -0.0044 | -0.703 | 2257 | 2277 |
| Armenian | Armenia_MBA | Greece_N | Mbuti | 0.0074 | 2.868 | 46368 | 45690 |
| Armenian | Armenia_MBA | Minoan_Lasithi | Mbuti | -0.0008 | -0.281 | 43117 | 43182 |
| Armenian | Armenia_MBA | Minoan_Odigitria | Mbuti | 0.0006 | 0.145 | 12181 | 12167 |
| Armenian | Armenia_MBA | Mycenaean | Mbuti | 0.0019 | 0.568 | 17154 | 17091 |
| Armenian | Armenia_MBA | Crete_Armenoi | Mbuti | -0.0049 | -0.567 | 2177 | 2199 |
| Armenian | Armenia_LBA | Greece_N | Mbuti | 0.0056 | 2.369 | 26917 | 26619 |
| Armenian | Armenia_LBA | Minoan_Lasithi | Mbuti | 0.0078 | **3.113** | 24936 | 24551 |
| Armenian | Armenia_LBA | Minoan_Odigitria | Mbuti | 0.0074 | 1.519 | 7062 | 6957 |
| Armenian | Armenia_LBA | Mycenaean | Mbuti | 0.0048 | 1.392 | 10136 | 10038 |
| Armenian | Armenia_LBA | Crete_Armenoi | Mbuti | 0.0114 | 1.041 | 1328 | 1298 |
| Armenian | Armenia_IA | Greece_N | Mbuti | 0.0013 | 0.518 | 51024 | 50892 |
| Armenian | Armenia_IA | Minoan_Lasithi | Mbuti | 0.004 | 1.482 | 45508 | 45146 |
| Armenian | Armenia_IA | Minoan_Odigitria | Mbuti | 0.0014 | 0.341 | 12037 | 12002 |
| Armenian | Armenia_IA | Mycenaean | Mbuti | -0.001 | -0.321 | 18437 | 18474 |
| Armenian | Armenia_IA | Crete_Armenoi | Mbuti | 0.0105 | 1.155 | 2113 | 2069 |
| Armenian | Armenia_ChL | Greek | Mbuti | 0.0001 | 0.073 | 27306 | 27299 |
| Armenian | Armenia_EBA | Greek | Mbuti | 0.0052 | 2.877 | 26818 | 26542 |
| Armenian | Armenia_MBA | Greek | Mbuti | 0.0018 | 0.808 | 24514 | 24425 |
| Armenian | Armenia_LBA | Greek | Mbuti | 0.0068 | **3.156** | 14612 | 14414 |
| Armenian | Armenia_IA | Greek | Mbuti | 0.003 | 1.334 | 26315 | 26155 |

**Supplementary Table 1.** The *D* statistics of the form (*D*(Modern_Armenians, Ancient_Armenian_Highland; *X_Balkan*, Mbuti)) based on SNP-chip data. Values with |Z| score > 3 are in bold.

| **Test population** | **t1** | **t2** |
|---|---|---|
| Chalcolithic Armenian samples | 1.00E-10 | 4.7E-02 |
| Early Bronze Age Armenian samples | 1.00E-10 | 2.9E-02 |
| Middle Bronze Age Armenian samples | 1.00E-10 | 9.5E-02 |
| Late Bronze Age Armenian samples | 1.00E-10 | 1.6E-01 |
| Iron Age Armenian samples | 1.00E-10 | 2.5E-02 |

**Supplementary Table 2.** Maximum likelihood estimated drift times in modern Armenian samples (t1) and ancient samples from the region (t2).

| **A** | **B** | **X** | **Y** | **D** | **Z score** | **ABBA** | BABA |
|---|---|---|---|---|---|---|---|
| Armenian | Armenia_ChL | Steppe_Eneolithic | Mbuti | -0.0209 | **-8.794** | 35717 | 37245 |
| Armenian | Armenia_EBA | Steppe_Eneolithic | Mbuti | -0.0091 | **-3.795** | 35569 | 36221 |
| Armenian | Armenia_MBA | Steppe_Eneolithic | Mbuti | -0.0178 | **-5.42** | 32693 | 33876 |
| Armenian | Armenia_LBA | Steppe_Eneolithic | Mbuti | -0.0131 | **-3.875** | 18596 | 19090 |
| Armenian | Armenia_IA | Steppe_Eneolithic | Mbuti | -0.0132 | **-4.29** | 33669 | 34568 |
| Armenian | Armenia_ChL | Steppe_EMBA | Mbuti | -0.0124 | **-7.502** | 51490 | 52784 |
| Armenian | Armenia_EBA | Steppe_EMBA | Mbuti | -0.0093 | **-5.31** | 50405 | 51355 |
| Armenian | Armenia_MBA | Steppe_EMBA | Mbuti | -0.0154 | **-6.629** | 45683 | 47112 |
| Armenian | Armenia_LBA | Steppe_EMBA | Mbuti | -0.0128 | **-6.238** | 26562 | 27252 |
| Armenian | Armenia_IA | Steppe_EMBA | Mbuti | -0.0132 | **-6.074** | 50601 | 51955 |
| Armenian | Armenia_ChL | Steppe_MLBA | Mbuti | -0.0116 | **-7.303** | 51516 | 52729 |
| Armenian | Armenia_EBA | Steppe_MLBA | Mbuti | -0.0064 | **-3.671** | 50512 | 51166 |
| Armenian | Armenia_MBA | Steppe_MLBA | Mbuti | -0.0103 | **-4.752** | 45871 | 46830 |
| Armenian | Armenia_LBA | Steppe_MLBA | Mbuti | -0.0078 | **-3.829** | 26689 | 27107 |
| Armenian | Armenia_IA | Steppe_MLBA | Mbuti | -0.0093 | **-4.376** | 50779 | 51733 |
| Armenian | Armenia_ChL | Steppe_IA | Mbuti | -0.0125 | **-4.898** | 42516 | 43593 |
| Armenian | Armenia_EBA | Steppe_IA | Mbuti | -0.0054 | -2.04 | 42047 | 42505 |
| Armenian | Armenia_MBA | Steppe_IA | Mbuti | -0.0115 | **-3.33** | 38636 | 39535 |
| Armenian | Armenia_LBA | Steppe_IA | Mbuti | -0.0019 | -0.562 | 22074 | 22158 |
| Armenian | Armenia_IA | Steppe_IA | Mbuti | -0.0101 | -2.882 | 40178 | 40997 |
| Armenian | Armenia_ChL | Anatolia_N | Mbuti | -0.0086 | **-5.317** | 51971 | 52870 |
| Armenian | Armenia_EBA | Anatolia_N | Mbuti | -0.0029 | -1.593 | 50979 | 51271 |
| Armenian | Armenia_MBA | Anatolia_N | Mbuti | 0.0036 | 1.612 | 46702 | 46370 |
| Armenian | Armenia_LBA | Anatolia_N | Mbuti | 0.0069 | **3.568** | 27198 | 26823 |
| Armenian | Armenia_IA | Anatolia_N | Mbuti | 0.0026 | 1.193 | 51671 | 51405 |

| | | | | | | | |
|---|---|---|---|---|---|---|---|
| Armenian | Armenia_ChL | Anatolia_BA | Mbuti | -0.0087 | **-3.351** | 32204 | 32767 |
| Armenian | Armenia_EBA | Anatolia_BA | Mbuti | -0.0066 | -2.453 | 31918 | 32341 |
| Armenian | Armenia_MBA | Anatolia_BA | Mbuti | 0.0019 | 0.551 | 30423 | 30306 |
| Armenian | Armenia_LBA | Anatolia_BA | Mbuti | -0.0005 | -0.159 | 17787 | 17807 |
| Armenian | Armenia_IA | Anatolia_BA | Mbuti | 0.0019 | 0.522 | 29938 | 29823 |
| Armenian | Armenia_ChL | Levant_N | Mbuti | -0.0086 | **-4.112** | 40467 | 41166 |
| Armenian | Armenia_EBA | Levant_N | Mbuti | -0.0056 | -2.656 | 39935 | 40386 |
| Armenian | Armenia_MBA | Levant_N | Mbuti | -0.0013 | -0.45 | 37285 | 37382 |
| Armenian | Armenia_LBA | Levant_N | Mbuti | 0.0068 | 2.406 | 21932 | 21638 |
| Armenian | Armenia_IA | Levant_N | Mbuti | 0.0013 | 0.425 | 38026 | 37930 |
| Armenian | Armenia_ChL | Levant_BA | Mbuti | -0.0103 | **-5.408** | 42396 | 43278 |
| Armenian | Armenia_EBA | Levant_BA | Mbuti | -0.0076 | **-3.772** | 41947 | 42590 |
| Armenian | Armenia_MBA | Levant_BA | Mbuti | -0.0005 | -0.194 | 39200 | 39241 |
| Armenian | Armenia_LBA | Levant_BA | Mbuti | 0.007 | 2.682 | 22928 | 22611 |
| Armenian | Armenia_IA | Levant_BA | Mbuti | 0.0038 | 1.407 | 39876 | 39572 |
| Armenian | Armenia_ChL | Iran_N | Mbuti | -0.0018 | -1.025 | 51168 | 51355 |
| Armenian | Armenia_EBA | Iran_N | Mbuti | -0.0075 | **-3.998** | 49738 | 50490 |
| Armenian | Armenia_MBA | Iran_N | Mbuti | -0.0063 | -2.665 | 45344 | 45916 |
| Armenian | Armenia_LBA | Iran_N | Mbuti | -0.0039 | -1.708 | 26366 | 26571 |
| Armenian | Armenia_IA | Iran_N | Mbuti | -0.0082 | **-3.416** | 50105 | 50929 |

**Supplementary Table 3.** The *D* statistics of the form (*D*(Modern_Armenians, Ancient_Armenian_Highland; *X_ancient_population*, Mbuti)) based on SNP-chip data. Values with |Z| score > 3 are in bold.

| **A** | **B** | **X** | **Y** | **D** | **Z score** | **ABBA** | **BABA** |
|---|---|---|---|---|---|---|---|
| Armenian | Armenia_ChL | Sardinian | Mbuti | 0.0015 | 0.91 | 27365 | 27283 |
| Armenian | Armenia_ChL | Sicilian | Mbuti | 0.0015 | 0.957 | 27254 | 27172 |
| Armenian | Armenia_ChL | Italian_North | Mbuti | -0.0003 | -0.184 | 27323 | 27339 |
| Armenian | Armenia_EBA | Sardinian | Mbuti | 0.0074 | **3.969** | 26894 | 26498 |
| Armenian | Armenia_EBA | Sicilian | Mbuti | 0.0066 | **3.625** | 26766 | 26416 |
| Armenian | Armenia_EBA | Italian_North | Mbuti | 0.0051 | 2.83 | 26841 | 26567 |
| Armenian | Armenia_MBA | Sardinian | Mbuti | 0.004 | 1.728 | 24587 | 24389 |
| Armenian | Armenia_MBA | Sicilian | Mbuti | 0.0032 | 1.398 | 24467 | 24313 |
| Armenian | Armenia_MBA | Italian_North | Mbuti | 0.0022 | 0.98 | 24545 | 24439 |
| Armenian | Armenia_LBA | Sardinian | Mbuti | 0.0106 | **4.716** | 14670 | 14362 |
| Armenian | Armenia_LBA | Sicilian | Mbuti | 0.0089 | **4.101** | 14590 | 14332 |
| Armenian | Armenia_LBA | Italian_North | Mbuti | 0.0068 | **3.133** | 14623 | 14424 |
| Armenian | Armenia_IA | Sardinian | Mbuti | 0.0056 | 2.392 | 26396 | 26102 |

| | | | | | | | |
|---|---|---|---|---|---|---|---|
| Armenian | Armenia_IA | Sicilian | Mbuti | 0.0046 | 2.002 | 26267 | 26027 |
| Armenian | Armenia_IA | Italian_North | Mbuti | 0.0026 | 1.154 | 26326 | 26192 |

**Supplementary Table 4.** The *D* statistics of the form (*D*(Modern_Armenians, Ancient_Armenian_Highland; *X_modern_population*, Mbuti)) based on SNP-chip data. Values with |Z| score > 3 are in bold.
